# Characterization of the microbiota associated with 12-week-old bovine fetuses exposed to divergent *in utero* nutrition

**DOI:** 10.1101/2021.09.14.460234

**Authors:** Samat Amat, Devin B. Holman, Kaycie Schmidt, Kacie L. McCarthy, Sheri T. Dorsam, Alison K. Ward, Pawel P. Borowicz, Lawrence P. Reynolds, Joel S. Caton, Kevin K. Sedivec, Carl R. Dahlen

## Abstract

A recent study reported the existence of a diverse microbiota in 5-to-7-month-old calf fetuses, suggesting that colonization of the bovine gut with so-called “pioneer” microbiota may begin during mid-gestation. In the present study, we investigated 1) the presence of microbiota in bovine fetuses at early gestation (12 weeks), and 2) whether the fetal microbiota is influenced by the maternal rate of gain or dietary supplementation with vitamins and minerals (VTM) during early gestation. Amniotic and allantoic fluids, and intestinal and placental (cotyledon) tissue samples obtained from fetuses (n = 33) on day 83 of gestation were processed for the assessment of fetal microbiota using 16S rRNA gene sequencing. The sequencing results revealed that a diverse and complex microbial community was present in each of these fetal compartments evaluated. Allantoic and amniotic fluids, and fetal intestinal and placenta microbiota each had distinctly different (0.047 ≥ R2 ≥ 0.019, P ≤ 0.031) microbial community structures. Allantoic fluid had a greater (P < 0.05) microbial richness (number of OTUs) (Mean 122) compared to amniotic fluid (84), intestine (63) and placenta (66). Microbial diversity (Shannon index) was similar for the intestinal and placental samples, and both were less diverse compared with fetal fluid microbiota (*P* < 0.05). Thirty-nine different archaeal and bacterial phyla were detected across all fetal samples, with Proteobacteria (55%), Firmicute*s* (16.2%), Acidobacteriota (13.6%) and Bacteroidota (5%) predominating. Among the 20 most relatively abundant bacterial genera, *Acidovorax, Acinetobacter, Brucella, Corynebacterium, Enterococcus, Exiguobacterium* and *Stenotrophomonas* differed by fetal sample type (*P* < 0.05). A total of 55 taxa were shared among the four different microbial communities. qPCR of bacteria in the intestine and placenta samples as well as scanning electron microscopy imaging of fetal fluids provided additional evidence for the presence of a microbiota in these samples. Minor effects of maternal rate of gain and VTM supplementation, and their interactions on microbial richness and composition were detected. Overall, the results of this study indicate that colonization with pioneer microbiota may occur during early gestation in bovine fetuses, and that that the maternal nutritional regime during gestation may influence the early fetal microbiota.

## INTRODUCTION

The neonatal gut harbors a low-diversity microbiota at birth; however, the gut microbial community undergoes developmental, transitional, and stable phases of progression before converging toward an adult-like state by the end of the first 3 to 5 years of life (Rodríguez et al., 2015; Stewart et al., 2018). Increasing evidence suggests that the early-life microbiome is important for the regulation of immune, endocrine, and metabolic developmental pathways (Robertson et al., 2019), and thus normal development of the early-life microbiome may be critical to health and wellbeing later in life (Arrieta et al., 2014; Tamburini et al., 2016). Many factors, including malnutrition, infections, and other lifestyle-related factors (e.g. mode of birth, breastfeeding, and antibiotic administration), can perturb normal assembly and evolution of the gut microbiome during infancy, resulting in long-lasting health complications (Willing et al., 2011; Vangay et al., 2015; Ranucci et al., 2017; Neuman et al., 2018; Shao et al., 2019). In addition to the factors that influence gut microbiome development during and after birth, emerging evidence of microorganisms in meconium (Nagpal et al., 2016; He et al., 2020), fetal fluids (Urushiyama et al., 2017; Stinson et al., 2019), and fetal intestine (Rackaityte et al., 2020; Mishra et al., 2021) suggests that microbial seeding of the infant intestine may begin *in utero*, and that factors influencing fetal microbial colonization may also shape postnatal microbiome development.

Although the *in utero* microbial colonization hypothesis is still controversial, and many support the “sterile-womb hypothesis” that infant microbiome acquisition occurs only during and after birth (Perez-Muñoz et al., 2017; de Goffau et al., 2019; Kennedy et al., 2021), very recent studies provide convincing evidence to support the former hypothesis. Rackaityte and colleagues (2020) were able to culture viable bacteria (*Micrococcaceae* and *Lactobacillus* strains) from the human fetal intestine at mid-gestation. Likewise, He et al. (2020) reported that seeding of the meconium microbiota is partially contributed by the microorganisms found in amniotic fluid. Furthermore, bacterial presence in the intestinal lumen of 14- and 18-week-old human (i.e., early second trimester) fetuses has recently been demonstrated by sequencing, imaging and culture-based approaches (Mishra et al., 2021). When coupled with emerging support for the potential role of the microbiome in the Developmental Origins of Health and Disease (DOHaD) (Stiemsma and Michels, 2018; Codagnone et al., 2019b), further research to evaluate the timing and mechanisms involved in the initial colonization of the fetal/infant gut is critical to a comprehensive understanding of development of the early-life gut microbiome (Chu et al., 2017; Walker et al., 2017).

Given that cattle have a similar singleton pregnancy and gestation period (280 days, 40 weeks) to humans, investigation of perinatal microbial colonization using a bovine animal model may provide more relevant information than other animals for clinical practices that are focused on the prevention of microbiome perturbations primarily during and/or after birth. Identifying the timeline for the emergence of the pioneer intestinal microbiota in cattle also has potential implications for harnessing the bovine microbiome to improve animal health and productivity. Until very recently, it was believed that colonization of the rumen by various microbes, including methanogenic archaea, begins as early as birth (Abecia et al., 2014; Guzman et al., 2015). However, this was challenged by recent evidence showing that prenatal microbial colonization may take place in the intestine of fetal calves. Results of a bovine caesarean section study that was conducted to evaluate bacterial load and the bacterial composition of the amniotic fluid and meconium of near full-term calves using 16S rRNA gene sequencing, qPCR and culturing suggested that in utero maternal-fetal bacterial transmission may occur before birth in calves (Husso et al., 2021). In addition, Guzman et al., (2020) investigated the presence of a microbiota in amniotic fluid and in 5 sample types from the gastrointestinal tract (GIT) of 5-, 6- and 7-month-old calf fetuses using both molecular and culture-based approaches. Using 16S rRNA gene sequencing, these authors showed that there are relatively diverse and distinct archaeal and bacterial communities present in the fetal GIT and amniotic fluid. Total bacterial abundance was also noted to increase with gestational age. The authors were also able to culture and isolate viable bacteria from these samples. This study, for the first time, provided sequencing and culture-based evidence that the intestine of calf fetuses is not sterile and that colonization by pioneer microbes may occur during mid-gestation.

In the present study, we investigated 1) the presence of microbiota in bovine fetuses at early gestation (12 weeks) using 16S rRNA gene sequencing, qPCR and SEM imaging; and 2) whether the fetal microbial composition is influenced by the maternal rate of gain and dietary vitamin and mineral (VTM) supplementation during early gestation. Of note, considering the potential involvement of the maternal microbiome in the DOHaD (Stiemsma and Michels, 2018; Calatayud et al., 2019; Codagnone et al., 2019a), and the potential subsequent impact of maternal microbiota alterations induced by dietary intervention on the development of the offspring microbiome (Calatayud et al., 2019), it is reasonable to hypothesize that maternal rate of gain would influence in utero maternal-fetal microbial cross-talk. A well-defined impact of maternal VTM supplementation exists on offspring health and performance in beef cattle with increased evidence highlighting the role of VTM in fetal programming during early gestation (Mee et al., 1995; Van Emon et al., 2020; Diniz et al., 2021; Menezes et al., 2021). Thus, we also were interested in investigating whether maternal VTM supplementation has an impact on perinatal microbial colonization.

## MATERIALS AND METHODS

All experimental procedures involving cattle were approved by the North Dakota State University Institutional Animal Care and Use Committee (protocol ID: A19012).

### Animal husbandry and experimental design

The fetuses were harvested from nulliparous Angus-cross heifers on d 83 ± 0.27 of gestation via ovariohysterectomy. The detailed information with respect to the experimental design, diet, feeding and animal husbandry has been described previously (Diniz et al., 2021; Menezes et al., 2021). Briefly, 35 angus-cross heifers (initial BW 359.5 ± 7.1 kg) were used in a randomized complete block design with a 2 × 2 factorial treatment arrangement. The main factors included vitamin and mineral supplementation (VTM vs. NoVTM), and the rate of gain (low gain [LG] vs. moderate gain [MG]). Heifers received a diet of 0.45 kg ground corn/heifer/day and 113 g/heifer/day of VTM premix (Purina Wind & Rain Storm All-Season 7.5 Complete, Land O’Lakes, Inc., Arden Hills, MN, USA) for at least 71 days before artificial insemination (AI) and continuing through the first 83 days of gestation. On the day of AI, within each VTM or NoVTM group, heifers were randomly assigned to one of two rate of gain groups: LG or MG. The LG group heifers were targeted to gain 0.28 kg/d during gestation and were fed a basal total mixed ration (consisting of triticale hay, corn silage, modified dried distillers’ grains plus solubles [DDGS] and ground corn). The MG group heifers were fed the basal total mixed ration plus the starch-based protein/energy supplement (a blend of ground corn, DDGS, wheat midds, fish oil, urea, and ethoxyquin) to achieve the targeted MG of 0.79 kg/d. Heifers receiving the VTM supplement (LG-VTM and MG-VTM) continued to receive the same concentration of VTM as during the pre-insemination period until the day of fetal collection. Heifers were bred by AI using female-sexed semen from a single sire. Pregnancy diagnosis was performed 35 days after AI, and fetal sex was determined on day 65 using transrectal ultrasonography. Only heifers confirmed pregnant with a female fetus continued on the experiment.

### Fetal fluids, and intestinal and placental tissue collections

On d 83 ± 0.27 of gestation, gravid reproductive tracts were collected via ovariohysterectomy as described previously (McLean et al., 2016). Immediately upon removal of the uterus, 20 ml of allantoic and amniotic fluid samples were collected using a sterile 20-mL Luer-lock syringe with a sterile18-auge needle as shown in Fig. 1S and were transferred into 50-mL Falcon tubes and immediately snap frozen using liquid nitrogen. In addition, the fetal intestine (Fig. 1S) was removed and one third portion of the intestine was dissected. The fetal (cotyledon) portion of the placenta was also dissected. Fetal intestinal and placental tissue samples were wrapped in aluminum foil and snap frozen. All samples were stored at −80°C until analyses.

**Figure 1.**
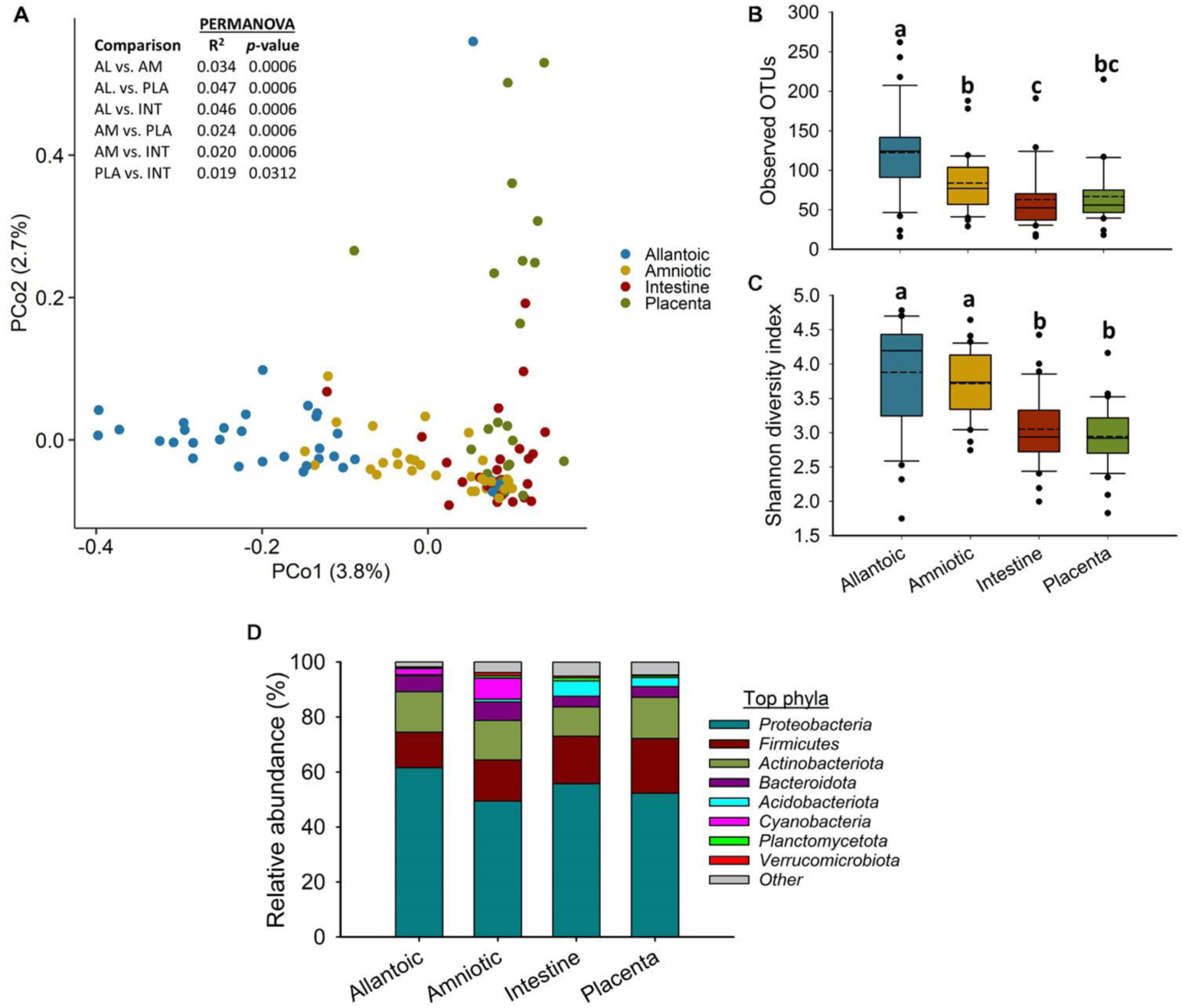
**A**) Principal coordinate analysis (PCoA) plot of the Bray-Curtis dissimilarities, **B**) the number of operational taxanomic units (OTUs), **C)** Shannon diversity indext and **D)** Percent relative abundance of the eight most relatively abundant phyla present in the allantoic and amniotic fluid and intestinal and placental microbiota of 83-day-old calf fetuses. Different lowercase letters indicate significantly different means (*P* < 0.05).

### Metagenomic DNA extraction

Metagenomic DNA from the allantoic and amniotic fluids, and fetal intestinal and placental tissue samples was extracted using cetyltrimethylammonium bromide buffer (CTAB) and a phenol:chloroform based method as described previously (Fujimura et al., 2016; Rackaityte et al., 2020) with some modifications. Ten milliliters of allantoic or amniotic fluid was centrifuged at maximum speed (16,000 × *g*) for 30 min at 4 °C using a floor ultra centrifuge (Beckman Optima L-60 Ultracentrifuge). After removal of the supernatant, the pellet was resuspended with 500 µl prewarmed CTAB buffer (10% CTAB, 0.7M NaCl, 240 mM potassium phosphate buffer pH 8), and transferred to a 2 mL Lysing Matrix E tube (containing 1.4 mm ceramic spheres, 0.1 mm silica spheres and one 4 mm glass bead) (MP Biomedical LLC, Irvine, CA, USA). The cell suspension was incubated in a ThermoMixer (Eppendorf, Enfield, CT) for 15 min at 65 °C and 800 rpm followed by mechanical cell lysis with bead beating (5.5 m/s, 30 s) in a FastPrep-24 (MP Biomedical LLC). Then, 500 µl of phenol:chloroform:isoamyl alcohol (25:24:1) was added into the cell lysate and mixed by manual shaking for 20 s before centrifugation at 16,000 × g for 5 min. The supernatant was carefully transferred into a heavy phase-lock gel tube (QuantaBio, Beverly, MA, USA), and mixed with an equal volume of chloroform and centrifuged at 16,000 × *g* for 5 min. The aqueous phase was then transferred into a sterile 1.5 mL tube and 1 µl linear acrylamide solution was added and vortexed. Then, 2 volumes of PEG-NaCl solution were added to the mixture and vortexed followed by incubation at 21 °C for 2 h. The isolated DNA was pelleted by centrifugation at 14,000 × g for 30 min and then washed twice with 70% ethanol. Finally, the isolated DNA was resuspended with 30 µl of 10 mM Tris-Cl pH 8, and the concentration was measured using a NanoDrop ND-1000 spectrophotometer and stored at −20 °C.

For DNA extraction from the intestinal and placental tissue samples, approximately 50 mg of tissue was dissected on a sterile empty Petri dish in a biosafety cabinet and placed in a 2 mL Lysing Matrix E tube. The remaining DNA extraction procedures were the same as described for fetal fluids except for an additional cycle of bead beating for the intestinal and placental tissue samples. DNA extraction from the nuclear free water (Corning Molecular Biology Grade Water, Corning, Glendale, AZ, USA) was performed to capture any potential microbial DNA contamination associated with the CTAB-based method.

### 16S rRNA gene sequencing and analysis

Amplification and sequencing of the 16S rRNA gene were performed at MR DNA (Shallowater, TX, USA) as described previously (Santiago-Rodriguez et al., 2016). Briefly, the V4 region of the 16S rRNA gene was amplified using the 515F (5’-GTGCCAGCMGCCGCGGTAA-3’) and 806R (5’-GGACTACHVGGGTWTCTAAT-3’) primers. A PCR clamp was also included to block amplification of bovine host DNA (Lundberg et al., 2013). PCR amplification was conducted using the HotStarTaq Plus Master Mix Kit (Qiagen Inc., Germantown, MD, USA). After amplification, PCR products were electrophoresed on a 2% agarose gel to ensure the correct size and band intensity. The 16S rRNA gene amplicons were indexed and pooled in equimolar concentrations and then purified using using calibrated Ampure XP beads. Then the pooled and purified PCR product was used to prepare an Illumina DNA library. These 16S rRNA gene libraries were then sequenced on an Illumina MiSeq instrument (Illumina Inc., San Diego, CA, USA) using the MiSeq reagent kit v3 (2 × 300 bp) following manufacturer’s instructions.

The 16S rRNA gene sequences were processed using DADA2 v. 1.18 (Callahan et al., 2016) in R. 4.0.3. Briefly, the forward and reverse reads were truncated at 210 bp, merged, chimeric sequences removed, and taxonomy assigned to each merged sequence, referred to here as operational taxonomic units (OTUs) at 100 % similarity, using the naïve Bayesian RDP classifier (Wang et al., 2007) and the SILVA SSU database release 138 (Quast et al., 2013). All OTUs that were found in the negative (extraction) control samples and likely to be contaminants were removed prior to analyses as were those OTUs classified as chloroplasts, eukaryote, or mitochondria. The number of OTUs per sample (richness), the Shannon and inverse Simpson’s diversity indices, and Bray-Curtis dissimilarities were calculated in R using Phyloseq 1.34.0 (McMurdie and Holmes, 2013) and vegan 2.5-7 (Oksanen et al., 2013). To account for uneven sequence depths, samples were randomly subsampled to 14,000 prior to the calculation of Bray-Curtis dissimilarities and diversity measures for comparisons among different sample types. For assessment of treatment effects within each sample type, the allantoic and amniotic fluid, and intestinal and placental tissue samples were randomly subsampled to 28,000, 20,000, 14,000 and 21,500 sequences, respectively. All 16S rRNA gene sequences are available in the NCBI’s sequence read archive under BioProject PRJNA731173.

### Bacterial concentration in fetal intestinal and placental tissue

The bacterial concentration was estimated for the fetal intestine and placental tissue samples using qPCR as previously described (Amat et al., 2020) with some modifications. Briefly, The primers 515Fq (5’-GTGYCAGCMGCCGCGGTAA-3’) (Parada et al., 2016) and 806Rq (5’-GGACTACNVGGGTWTCTAAT-3’ (Apprill et al., 2015) were used to amplify the V4 region of the 16S rRNA gene. Each qPCR mixture contained 1 SsoAdvanced Universal SYBR Green Supermix (Bio-Rad Laboratories, Inc. Hercules, California), 0.4 µM (each) primer, 0.1 µg/µl bovine serum albumin (New England Biolabs, Pickering, ON, Canada), and 30 ng of DNA, in a total volume of 25 µl. A CFX96 Touch Real-Time PCR Detection system (Bio-Rad Laboratories Ltd) was used with the following conditions: an initial denaturation at 95 °C for 3 min, followed by 40 cycles at 95 °C for 25 s, 50 °C for 30 s, and then 72 °C for 45 s. Standard curves (10^2^ to 10^8^ gene copies) were generated using the pDrive cloning vector (Qiagen Inc.) containing the PCR product from 16S rRNA gene. All qPCRs were performed in duplicate with standards (1 µl) and no-template controls (1 µl of nuclease-free H_2_O), as well as a positive control (DNA from bovine deep nasopharyngeal swab). A melt curve analysis was performed following qPCR amplification to ensure that only target genes were amplified. The copy number was normalized by 30 ng of input DNA.

### Scanning electron microscopy imaging

Scanning electron microscopy (SEM) imaging was performed on some of the allantoic and amniotic fluid samples that had greatest DNA concentrations. The SEM imaging was carried out by the NDSU Electron Microscopy Center core facility. Allantoic and amniotic fluids (4 ml) were centrifuged for 10 minutes at 14,000 × *g* to produce a pellet, then the pellet was resuspended in deionized water. This process was repeated twice to remove buffer salts. Pelleted material was then applied to round glass coverslips affixed to aluminum mounts with silver paint (SPI Supplies, West Chester PA, USA), air dried, and sputter coated with carbon (Cressington 208c, Ted Pella Inc., Redding, CA, USA). Images were obtained using a JEOL JSM-7600F scanning electron microscope at an accelerating voltage of 2 kV or a JEOL JSM-6490LV scanning electron microscopy at an accelerating voltage of 15 kV (JEOL USA Inc., Peabody, MA, USA).

### Statistical analysis

Differentially abundant genera within the 20 most relatively abundant genera among the four sample types were identified using an ANOVA with sample type and treatment in the model. The Benjamini-Hochberg procedure was used to correct all *P*-values for multiple comparisons. The effect of fetal sample type and the treatment within each sample type on the microbial community structure was assessed using Bray-Curtis dissimilarities and PERMANOVA (adonis2 function). Pairwise comparisons of the Bray-Curtis dissimilarities among sample types and treatment groups were done using the R package pairwise Adonis v. 0.01 with the Benjamini-Hochberg procedure used to correct *P*-values for multiple comparisons.

The number of OTUs (richness), diversity indices, relative abundance of the most relatively abundant phyla and genera between sample types (allantoic and amniotic fluids, and intestinal and placental tissues) were compared using the generalized liner mixed model estimation procedure (PROC GLIMMIX) in SAS (ver. 9.4, SAS Institute Inc. Cary, NC). The data regarding the comparison among different dietary treatment groups within each sample type were analyzed as 2 × 2 factorial treatment arrangement using the PROC GLIMMIX. The model included main effects of VTM supplementation (VTM or NoVTM), rate of gain (LG or MG) and the respective interaction. Means among different sample types or treatment groups within each sample type were compared using the LSMEANS statement and significance was declared at *P* < 0.05.

### RESULTS

### Sequencing Results

An average of 105,458 ± 77,752 (SD) 16S rRNA gene sequences per sample (min. = 14,029; max. = 514,084) were obtained from 130 allantoic and amniotic fluid, and intestinal and placental tissue samples from the 83-day-old calf fetuses. From these sequences, a total of 8,544 archaeal and bacterial OTUs (at 100% sequence similarity) were identified and classified into 39 unique phyla (3 archaeal and 36 bacterial phyla), and 921 unique genera.

### Microbial community composition and structure of the allantoic and amniotic fluid, and intestinal and placental tissues

A relatively diverse and unique microbial community was detected in the four fetal sample types (Fig.1). The microbial community structure was significantly different between the allantoic and amniotic fluid, and intestinal and placental tissue-associated microbiota (0.047 ≥ R^2^ ≥ 0.019, *P* ≤ 0.031) (Fig. 1A). Alpha diversity metrics also differed by fetal sample type (Fig 1B and C). Allantoic fluid had a greater (*P* < 0.05) microbial richness (number of OTUs, at 97% sequence similarity) (122 ± 10) compared with amniotic fluid (84 ± 6) and the intestinal (63 ± 7) and placental (66 ± 6) tissue. The number of OTUs in the amniotic fluid was similar to the placenta (*P* > 0.05) but greater than in the intestine (*P* < 0.05). Microbial diversity (Shannon index) was similar for the intestinal and placental samples, and both were less diverse compared to the fetal fluid (*P* < 0.05).

Among the 39 different phyla detected across all fetal samples, Proteobacteria (54.8%) was the most relatively abundant phylum followed by Firmicutes (16.3%), Actinobacteriota (13.7%), Bacteroidota (5.0%), and Acidobacteriota (2.6%) (Fig. 1D). Archaeal phyla (Crenarchaeota, Euryarchaeota and Nanoarchaeota) accounted for only 0.34% of the 16S rRNA gene sequences (Table S1). The relative abundance of some of the bacterial phyla also differed by fetal sample type (Fig. 1D). Proteobacteria was less relatively abundant in amniotic fluid (49.4%) compared to allantoic fluid (61.6%), and intestinal (55.7%) and placental (53.6%) tissues (*P* < 0.05). The fetal intestine (5.5%) and placenta (3.2%) harbored a greater relative abundance of *Acidobacteriota* compared to the allantoic (0.5%) and amniotic (1.0%) fluids (*P* < 0.05).

At the genus level, *Acidovorax, Acinetobacter, Stenotrophomonas, Brucella, Anoxybacillus, Sphingomonas, Allorhizobium-Neorhizobium-Pararhizobium-Rhizobium* (ANPR), *Salinisphaera* and *Lactobacillus* were the most relatively abundant genera among all samples (Fig. 2). Although between sample variance was relatively high, *Acidovorax, Acinetobacter, Brucella, Corynebacterium, Enterococcus, Exiguobacterium* and *Stenotrophomonas* were differentially abundant between different sample types (*P* < 0.05). *Acidovorax* was equally abundant across amniotic fluid, and intestinal and placental tissues (*P* > 0.05) but less abundant in allantoic fluid (*P* < 0.05). Placental samples had a greater relative abundance of *Acinetobacter* compared to other sample types. ANPR and *Brucella* were more relatively abundant in allantoic fluid, whereas *Salinisphaera* and *Aphanizomenon* were enriched in amniotic fluid (*P* < 0.05).

**Figure 2.**
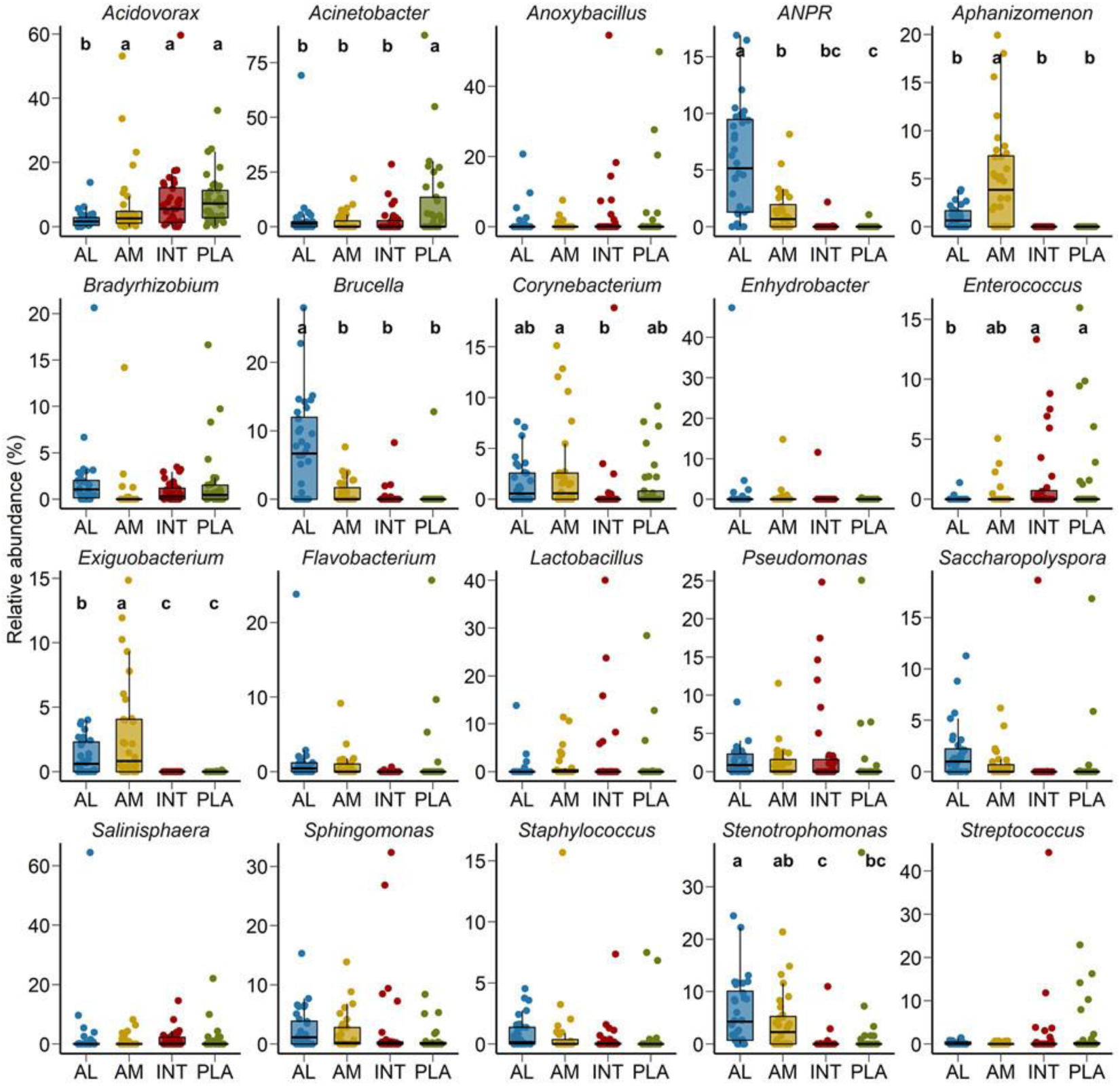
**Percent** relative abundance of the 20 most relatively abundant bacterial genera in the allantoic (AL) and amniotic (AM) fluid, and intestinal (INT) and placental (PLA) tissue microbiota in 83-day-old calf fetuses.

### Shared taxa in the allantoic and amniotic fluid, and intestinal and placental microbiota

No single OTU was identified in 50% of all the allantoic and amniotic fluid, and intestinal and placenta samples. As shown in the heatmap of the 100 most abundant OTUs (Fig. 3), there was considerable inter-individual variation in both prevalence and abundance of most of these taxa. Certain OTUs including OTU96 (*Pseudomonas*), OTU22 (*Acinetobacter)*, OTU78 (*Achromobacter*), OTU18 (*Brucella*), OTU26 (ANPR), OTU25 (*Caulobacteraceae*) and OTU47 (*Saccharopolyspora*) were present in the majority of the allantoic fluid samples with greater abundance. From these OTUs, OTU18 (*Brucella*), OTU26 (ANPR) and OTU47 (*Saccharopolyspora*) were also more prevalent in the amniotic fluid samples. Relatively more prevalent taxa detected from fetal intestinal samples are OTU17 (*Anoxybacillus)* and OTU70 (*Microbacteriaceae*). In placental tissue, two taxa (OTU19 and OTU14) within the genus *Acinetobacter* were more frequently detected.

**Figure 3.**
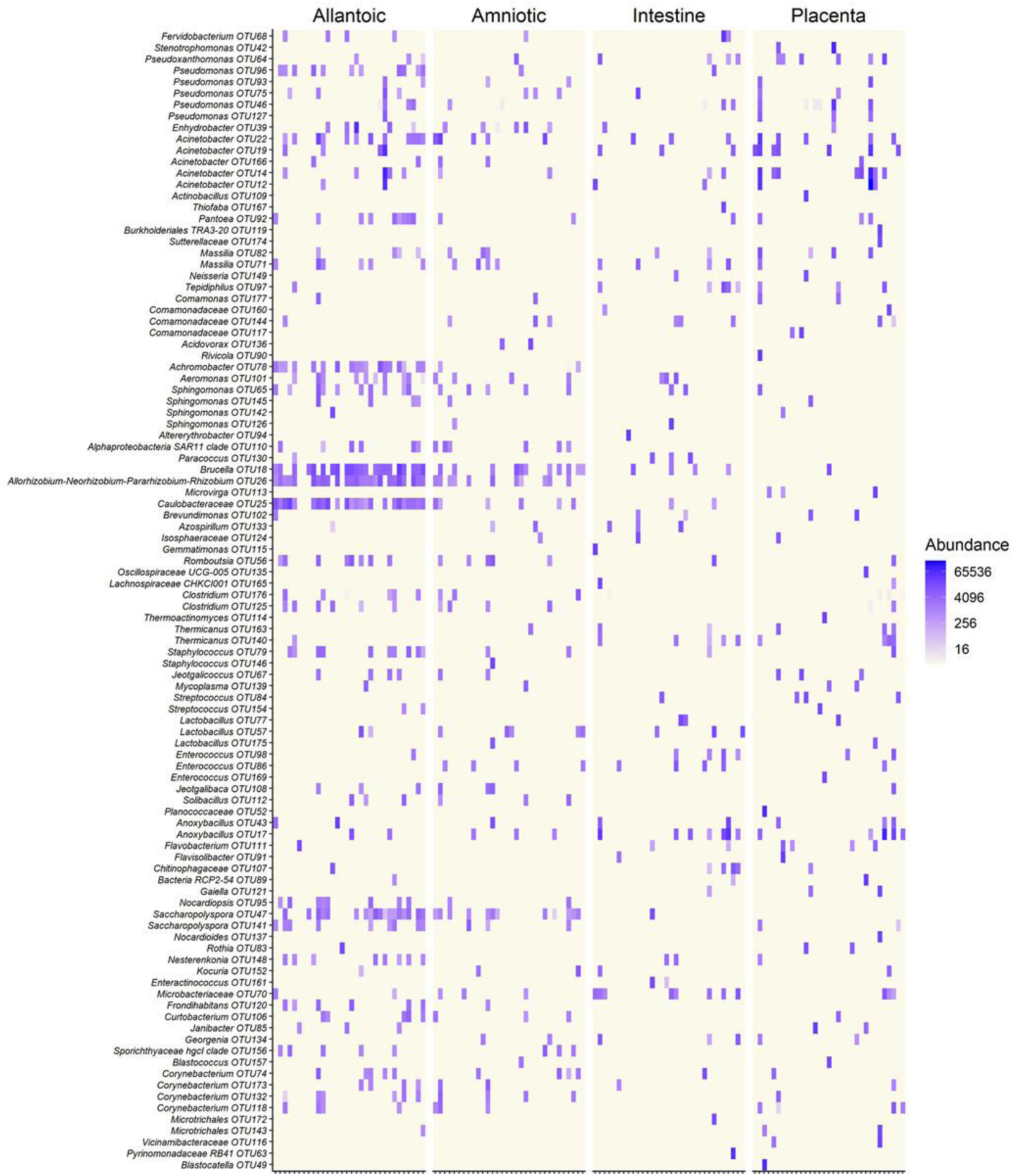
Heatmap showing the 100 most abundant OTUs (log_4_ transformed) within the allantoic and amniotic fluid, and intestinal and placental microbiota in 83-day-old calf fetuses.

A Venn diagram displaying the number of unique and shared OTUs among different sample types is displayed in Fig. 4. Overall, the proportion of unique OTUs identified in amniotic and allantoic fluid, and intestinal and placental microbiota accounted for 33%, 23%, 16% and 17% of the OTUs, respectively. A total of 223 (3%) OTUs were shared by allantoic and amniotic fluids, and 2% of the OTUs were shared between intestine and placental tissues. Less than 1% taxa were shared between fetal fluid and tissue samples or between the three of the four sample types. Only 55 OTUs (< 1% of the total OTUs) were shared among the four different microbial communities.

**Figure 4.**
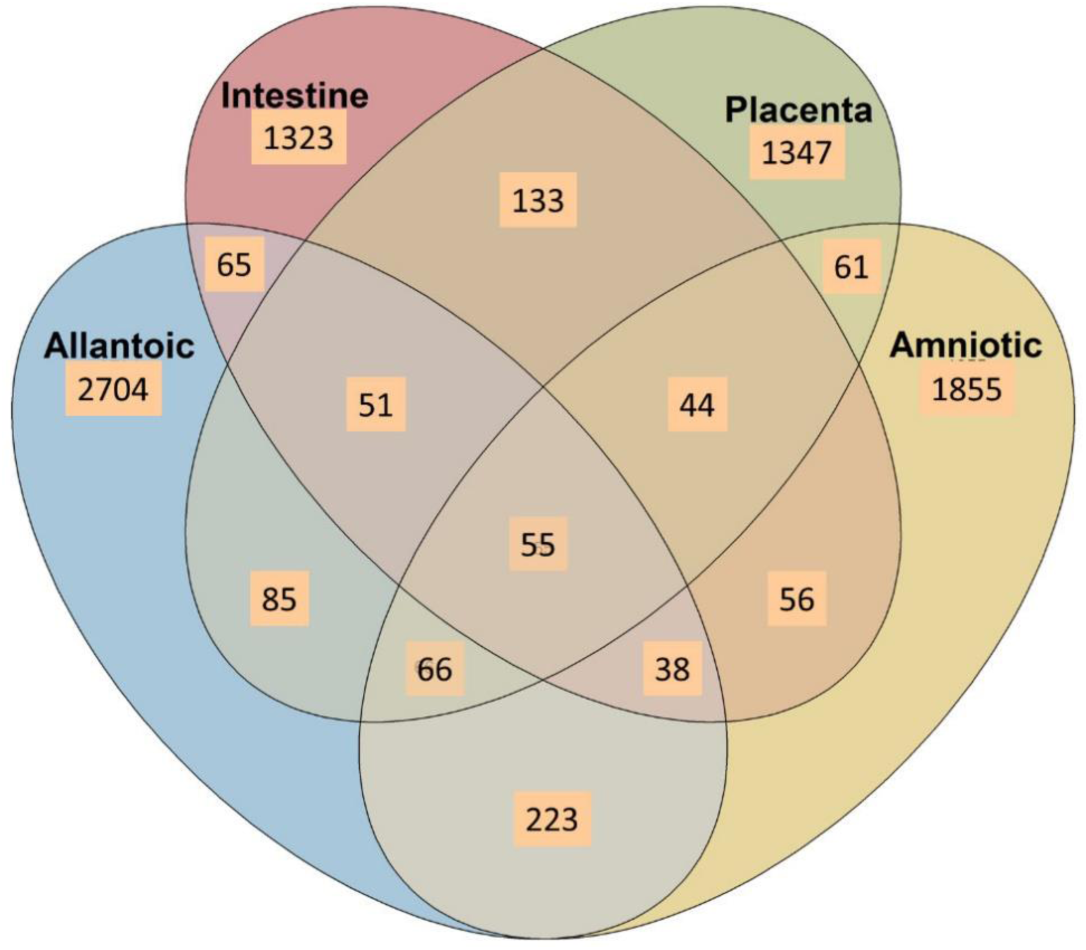
Venn diagram displaying the number of shared and unique OTUs among the allantoic and amniotic fluid and intestinal and placental microbiota in 83-day-old calf fetuses.

### Quantitative PCR and scanning electron microscopy imaging-based evidence supports the presence of microbiota in bovine fetuses

To further confirm the 16S rRNA gene sequencing results we quantified the total bacterial concentration in the fetal intestinal and placental tissue samples using qPCR. The total bacterial concentration in placental tissue samples was estimated to be 46% greater than that of intestinal tissue (*P* < 0.0001). Next, to provide visual evidence of the presence of microbial cells in the fetal fluid samples, SEM imaging was performed on select amniotic (n = 2) and allantoic fluid (n =3) samples. Microbial cells with prokaryotic cell structure (cocci, spherical) were detected in two allantoic fluid samples (Fig. 5).

**Figure 5.**
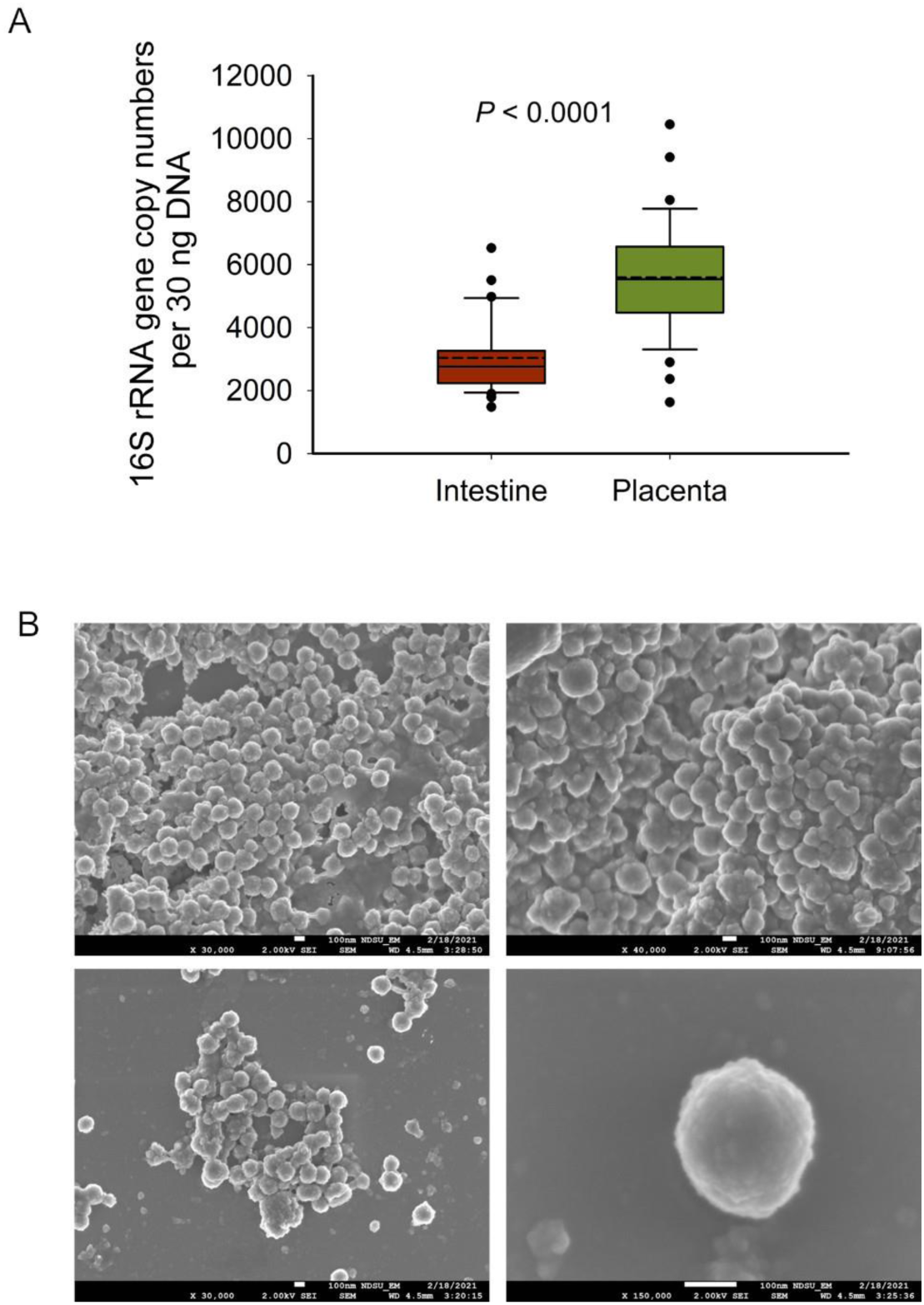
(**A**) The total bacterial abundance in intestinal and placental tissue samples estimated by qPCR and (**B**) scanning electron microscopy (SEM) images of microbial cells detected from fetal fluid samples of 83-day-old calf fetuses; white bars at the bottom of each image indicate 100 nm.

### The effect of maternal rate of gain and VTM supplementation on the fetal microbiota

We also evaluated whether the fetal microbiota in early gestation would diverge in response to *in utero* environment resulting from altered maternal nutrition. To this end, we analyzed the sequencing data by maternal nutritional treatment groups within each fetal sample type. Microbial community structure did not differ between treatment groups in any of amniotic and allantoic fluid, and intestinal and placental microbiota in 83-day-old bovine fetuses (0.0152 ≥ R^2^ ≥ 0.0762, *P* ≥ 0.18) (Fig. 6).

**Figure 6.**
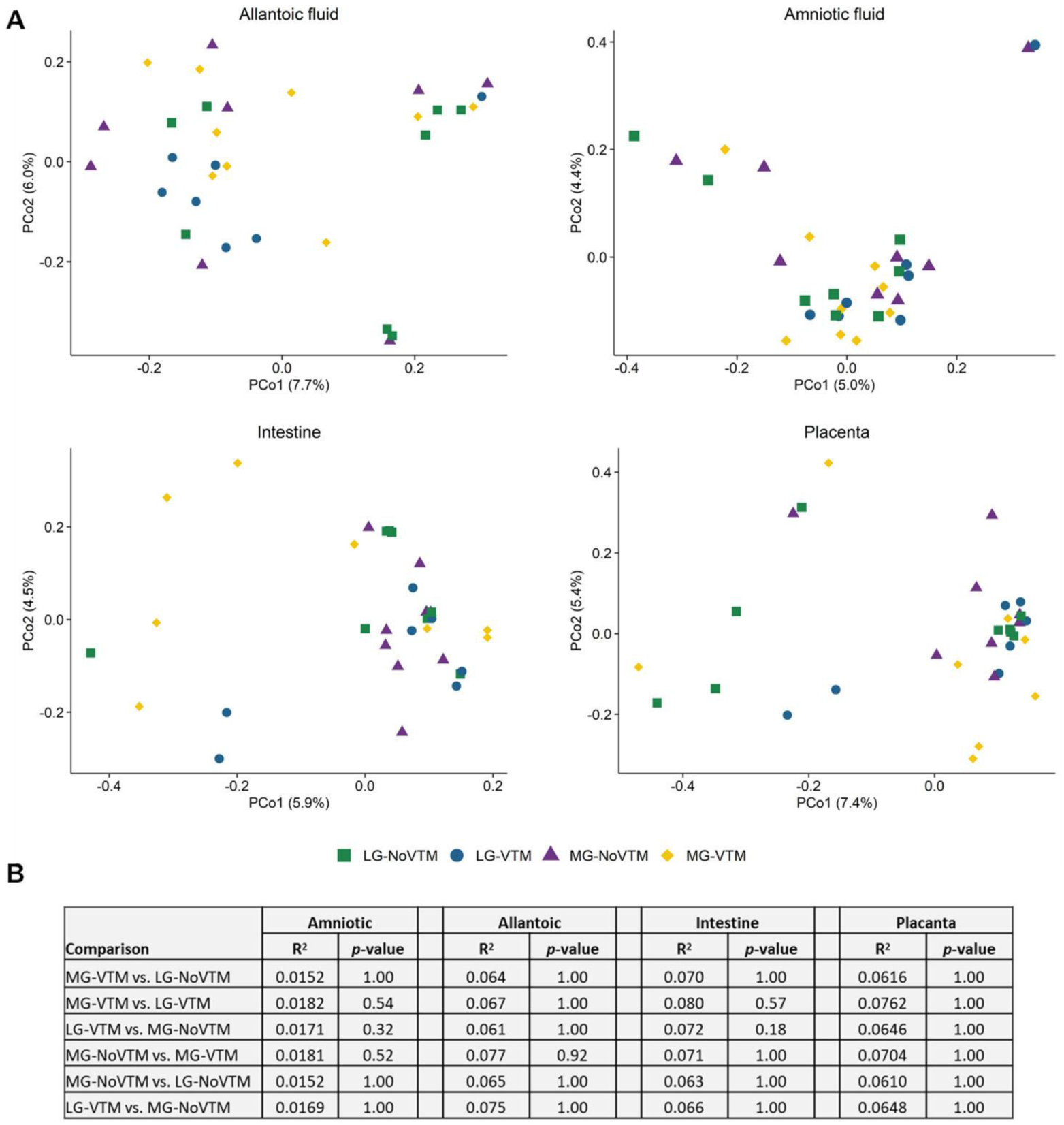
**(A)** Principal coordinate analysis (PCoA) plots of the Bray-Curtis dissimilarities by maternal nutrition treatment in amniotic and allantoic fluids, and intestinal and placental tissue microbiota of 83-day-old calf fetuses. The percentage of variation explained by each principal coordinate is indicated on the axes; (**B**) summary of the PERMANOVA comparisons of the Bray-Curtis dissimilarities among maternal treatment groups within each sample type. The fetuses were harvested from of dams that received one of four diets: no vitamin and mineral supplementation and targeted to moderate gain (MG-NoVTM); no vitamin and mineral supplementation and targeted to low gain (LG-NoVTM); vitamin and mineral supplementation and targeted to moderate gain (MG-VTM); or vitamin and mineral supplementation and targeted to low gain (LG-VTM).

No effects of VTM supplementation, rate of gain or their respective interaction were observed for microbial richness (number of OTUs) of amniotic fluid- and placenta-associated microbiota (*P* ≥ 0.122) (Fig. 7, Table S2). However, the number of OTUs observed in allantoic fluid samples derived from the fetuses born from the LG-NoVTM heifers was greater than that of MG-NoVTM fetuses (*P* < 0.05). The interaction effect was significant for the microbial richness of intestinal microbiota (*P* = 0.05), with MG-VTM fetuses had greater microbial richness in their intestine compared to MG-NoVTM fetuses (P < 0.05). Microbial diversity (Shannon and Inverse Simpson’s diversity indices) was not impacted by main effects of VTM, rate of gain, or their respective interactions in any of the fetal sample type (*P* > 0.10).

**Figure 7.**
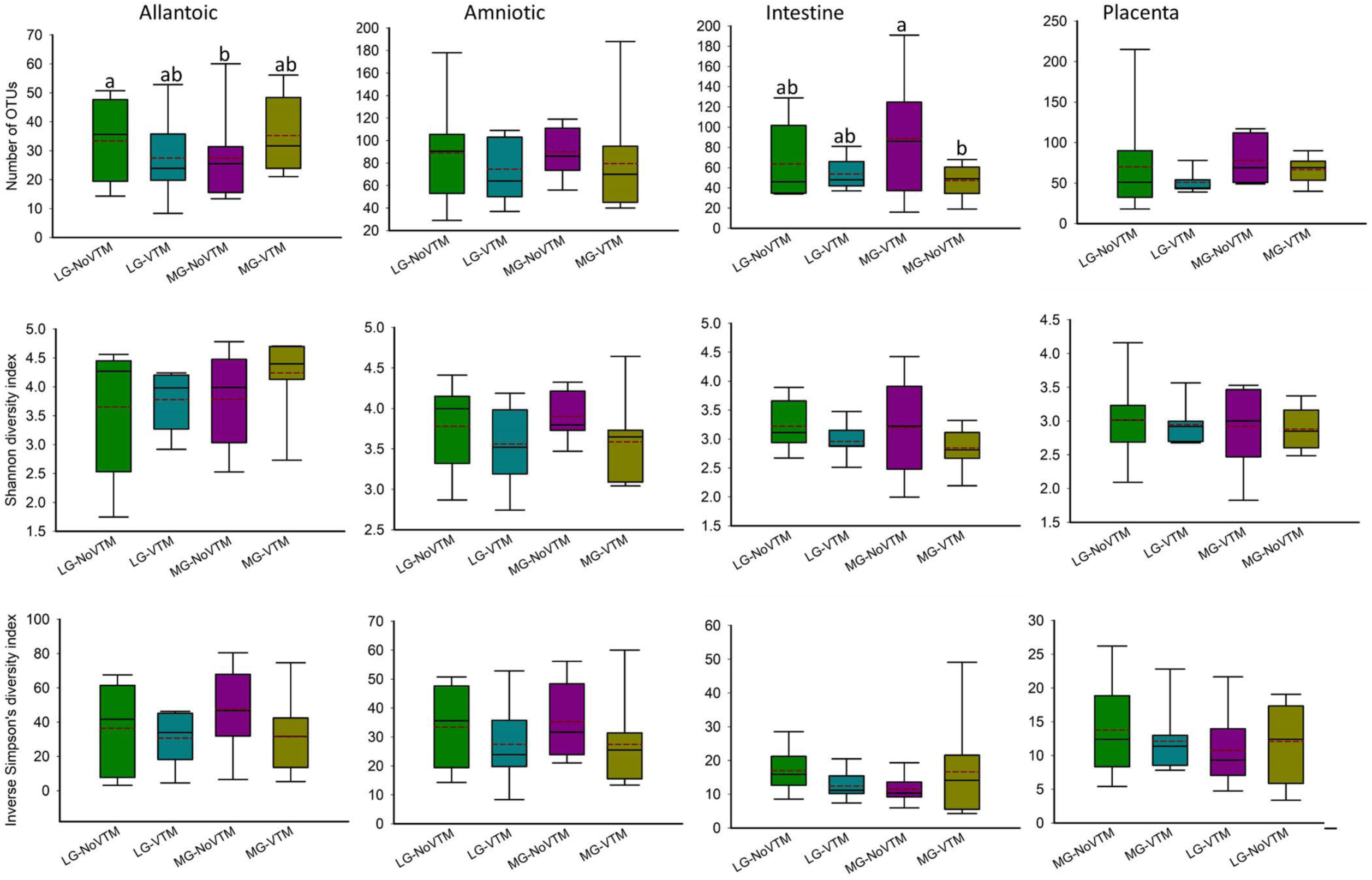
The effects of rate of gain, and vitamin and mineral supplementation on alpha diversity metrics of the allantoic and amniotic fluid, and intestinal and placental microbiota in 83-day-old calf fetuses (n = 33). The different lowercase letters indicate significantly different means (*P* < 0.05). LG-noVTM = low gain, no vitamin-mineral supplement; LG-VTM = low gain, vitamin-mineral supplement; MG-No VTM = moderate gain, no vitamin-mineral supplement; MG-VTM = moderate gain, vitamin-mineral supplement.

When the microbial composition was compared at the phylum level, no significant differences were noted between treatment groups for any of the five most relatively abundant phyla in allantoic fluid (*P* > 0.05) (Fig. 8, Table S2). However, the relative abundance of some of the five most relatively abundant phyla in amniotic, and intestinal and placental tissues was affected by treatment (VTM supplement or rate of gain or the interactions). For example, the effect of gain and VTM interaction on Proteobacteria in both amniotic fluid and intestinal tissue was significant (*P* ≤ 0.04). The relative abundance of Cyanobacteria in amniotic fluid and Firmicutes in intestine was affected by VTM supplementation (*P* ≤ 0.014). Rate of gain tended to affect the relative abundance of Bacteroidota (*P* = 0.057) in amniotic fluid and Firmicutes (*P* = 0.066) in placental tissue.

**Figure 8.**
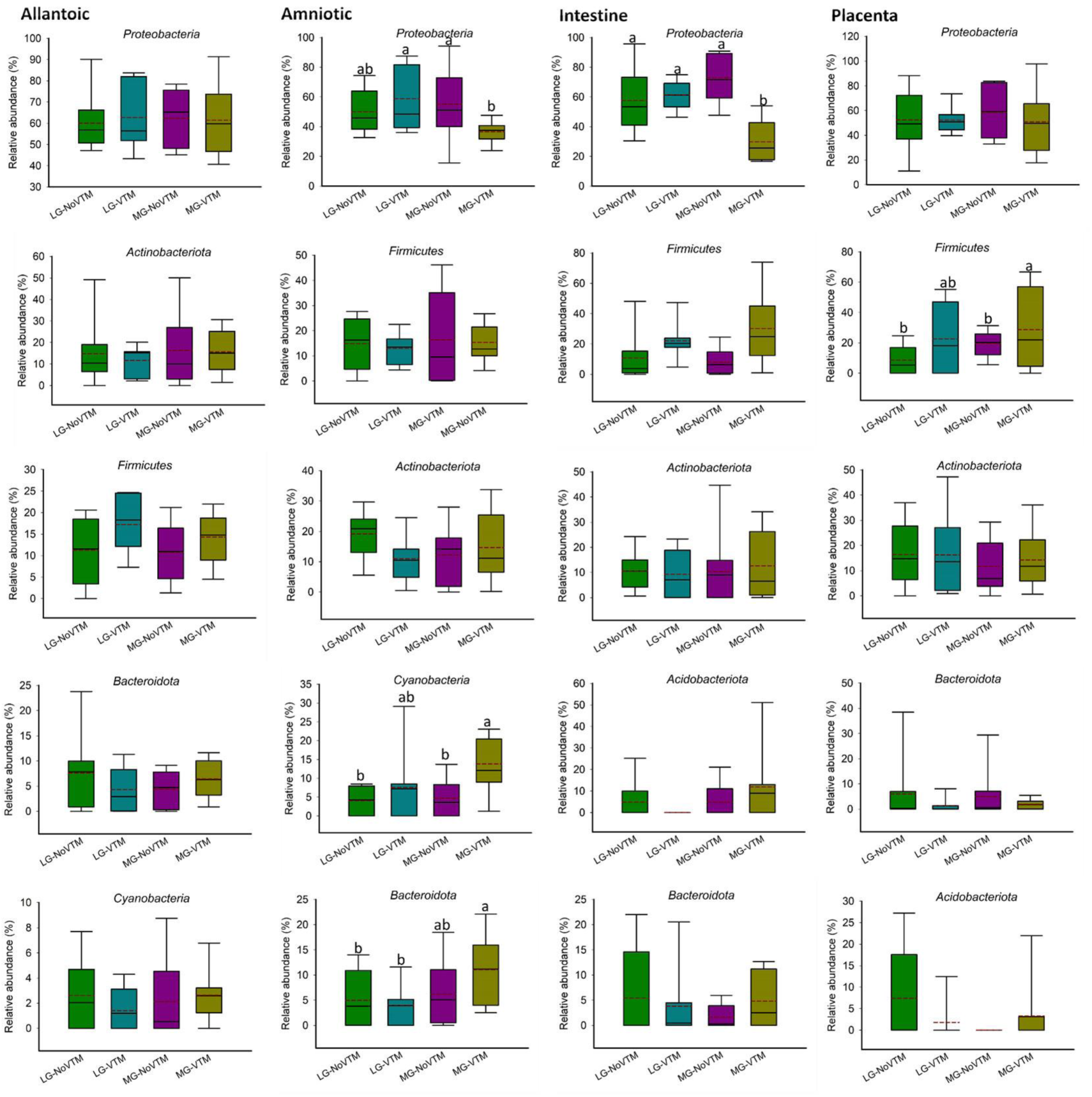
The effects of rate of gain, and vitamin and mineral supplementation on the relative abundance of the five most relatively abundant phyla present in allantoic and amniotic fluid, and intestine and placental microbiota in 83-day-old calf fetuses. Different lowercase letters indicate significantly different means (*P* < 0.05).

At the genus level, among the eight most relatively abundant genera listed in Table 1, the relative abundance of *Sphingomonas* in allantoic fluid, *Salinisphaera* in the intestine and *Streptococcus* in the placenta was affected by the rate of gain (*P* ≤ 0.037) (Table 1). A significant effect of VTM supplementation was observed on the relative abundance of *Aphanizomenon* (*P* = 0.010). The effect of VTM supplementation (*P* = 0.072), and an interaction between VTM and rate of gain (*P* = 0.067) tended to be significant for the relative abundance of *Anoxybacillus* in placenta. The relative abundance of the remaining eight most predominant genera was not influenced by the treatment (*P* > 0.10).

**Table 1.**
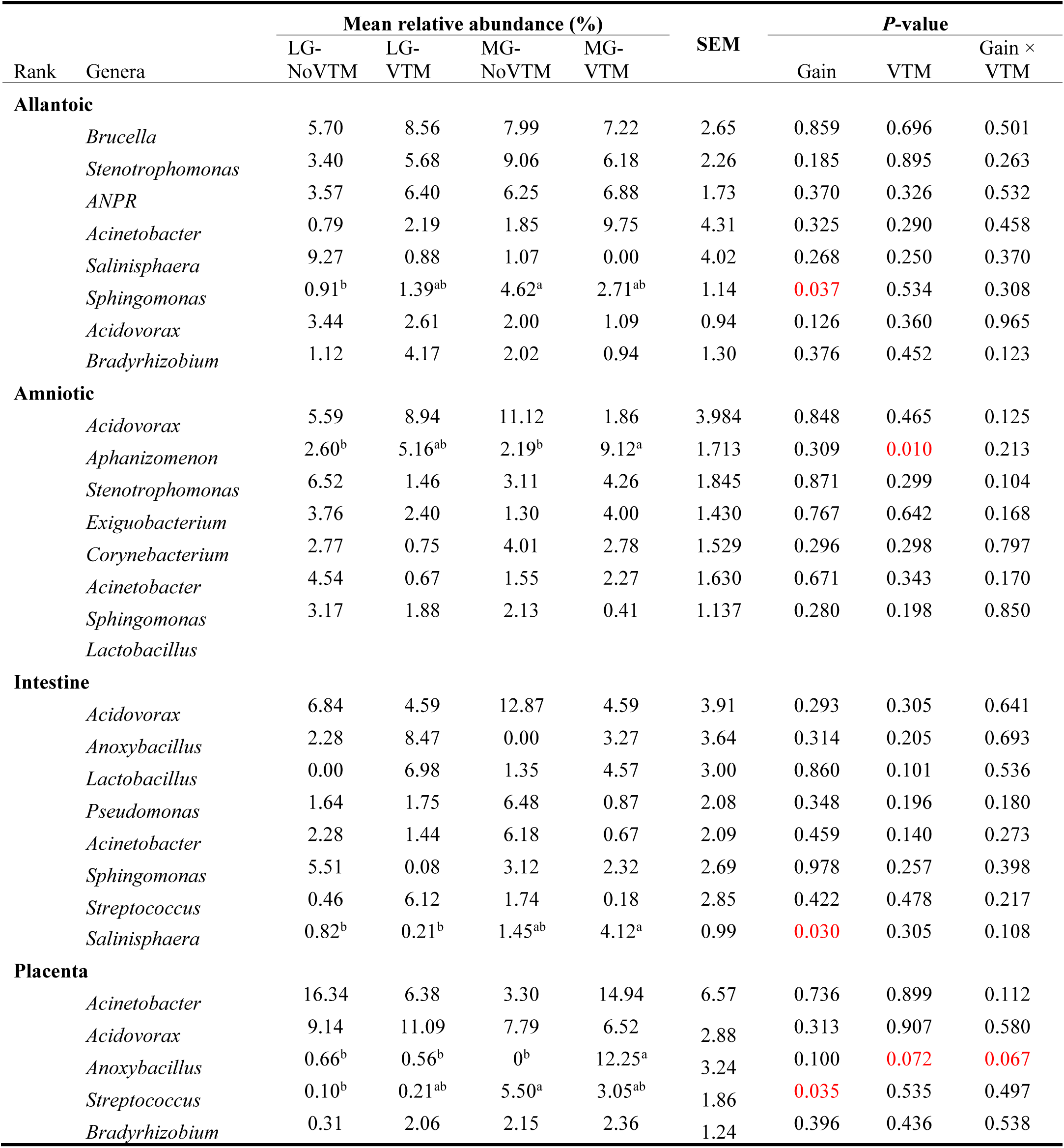
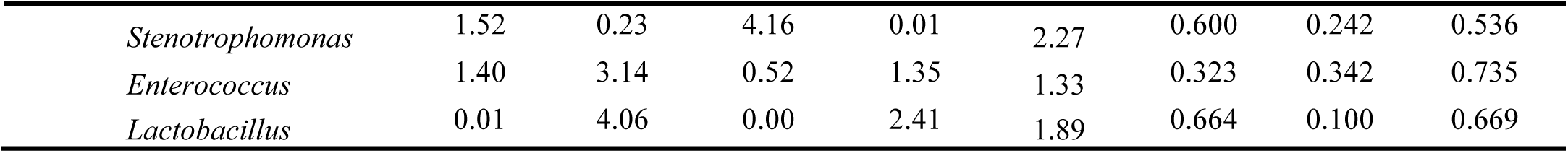
The effects of rate of gain, and vitamin and mineral supplementation on the relative abundance of the 8 most relatively abundant genera present in allantoic and amniotic fluid, and intestine and placental microbiota in 83-day-old calf fetuses.

| Rank | Genera | Mean relative abundance (%) |  |  |  | SEM | P-value |  |  |
| --- | --- | --- | --- | --- | --- | --- | --- | --- | --- |
|  |  | LG-NoVTM | LG-VTM | MG-NoVTM | MG-VTM |  | Gain | VTM | Gain ×<br>VTM |
| Allantoic |  |  |  |  |  |  |  |  |  |
|  | *Brucella* | 5.70 | 8.56 | 7.99 | 7.22 | 2.65 | 0.859 | 0.696 | 0.501 |
|  | *Stenotrophomonas* | 3.40 | 5.68 | 9.06 | 6.18 | 2.26 | 0.185 | 0.895 | 0.263 |
|  | *ANPR* | 3.57 | 6.40 | 6.25 | 6.88 | 1.73 | 0.370 | 0.326 | 0.532 |
|  | *Acinetobacter* | 0.79 | 2.19 | 1.85 | 9.75 | 4.31 | 0.325 | 0.290 | 0.458 |
|  | *Salinisphaera* | 9.27 | 0.88 | 1.07 | 0.00 | 4.02 | 0.268 | 0.250 | 0.370 |
|  | *Sphingomonas* | 0.91<sup>b</sup> | 1.39<sup>ab</sup> | 4.62<sup>a</sup> | 2.71<sup>ab</sup> | 1.14 | 0.037 | 0.534 | 0.308 |
|  | *Acidovorax* | 3.44 | 2.61 | 2.00 | 1.09 | 0.94 | 0.126 | 0.360 | 0.965 |
|  | *Bradyrhizobium* | 1.12 | 4.17 | 2.02 | 0.94 | 1.30 | 0.376 | 0.452 | 0.123 |
| Amniotic |  |  |  |  |  |  |  |  |  |
|  | *Acidovorax* | 5.59 | 8.94 | 11.12 | 1.86 | 3.984 | 0.848 | 0.465 | 0.125 |
|  | *Aphanizomenon* | 2.60<sup>b</sup> | 5.16<sup>ab</sup> | 2.19<sup>b</sup> | 9.12<sup>a</sup> | 1.713 | 0.309 | 0.010 | 0.213 |
|  | *Stenotrophomonas* | 6.52 | 1.46 | 3.11 | 4.26 | 1.845 | 0.871 | 0.299 | 0.104 |
|  | *Exiguobacterium* | 3.76 | 2.40 | 1.30 | 4.00 | 1.430 | 0.767 | 0.642 | 0.168 |
|  | *Corynebacterium* | 2.77 | 0.75 | 4.01 | 2.78 | 1.529 | 0.296 | 0.298 | 0.797 |
|  | *Acinetobacter* | 4.54 | 0.67 | 1.55 | 2.27 | 1.630 | 0.671 | 0.343 | 0.170 |
|  | *Sphingomonas* | 3.17 | 1.88 | 2.13 | 0.41 | 1.137 | 0.280 | 0.198 | 0.850 |
|  | *Lactobacillus* |  |  |  |  |  |  |  |  |
| Intestine |  |  |  |  |  |  |  |  |  |
|  | *Acidovorax* | 6.84 | 4.59 | 12.87 | 4.59 | 3.91 | 0.293 | 0.305 | 0.641 |
|  | *Anoxybacillus* | 2.28 | 8.47 | 0.00 | 3.27 | 3.64 | 0.314 | 0.205 | 0.693 |
|  | *Lactobacillus* | 0.00 | 6.98 | 1.35 | 4.57 | 3.00 | 0.860 | 0.101 | 0.536 |
|  | *Pseudomonas* | 1.64 | 1.75 | 6.48 | 0.87 | 2.08 | 0.348 | 0.196 | 0.180 |
|  | *Acinetobacter* | 2.28 | 1.44 | 6.18 | 0.67 | 2.09 | 0.459 | 0.140 | 0.273 |
|  | *Sphingomonas* | 5.51 | 0.08 | 3.12 | 2.32 | 2.69 | 0.978 | 0.257 | 0.398 |
|  | *Streptococcus* | 0.46 | 6.12 | 1.74 | 0.18 | 2.85 | 0.422 | 0.478 | 0.217 |
|  | *Salinisphaera* | 0.82<sup>b</sup> | 0.21<sup>b</sup> | 1.45<sup>ab</sup> | 4.12<sup>a</sup> | 0.99 | 0.030 | 0.305 | 0.108 |
| Placenta |  |  |  |  |  |  |  |  |  |
|  | *Acinetobacter* | 16.34 | 6.38 | 3.30 | 14.94 | 6.57 | 0.736 | 0.899 | 0.112 |
|  | *Acidovorax* | 9.14 | 11.09 | 7.79 | 6.52 | 2.88 | 0.313 | 0.907 | 0.580 |
|  | *Anoxybacillus* | 0.66<sup>b</sup> | 0.56<sup>b</sup> | 0<sup>b</sup> | 12.25<sup>a</sup> | 3.24 | 0.100 | 0.072 | 0.067 |
|  | *Streptococcus* | 0.10<sup>b</sup> | 0.21<sup>ab</sup> | 5.50<sup>a</sup> | 3.05<sup>ab</sup> | 1.86 | 0.035 | 0.535 | 0.497 |
|  | *Bradyrhizobium* | 0.31 | 2.06 | 2.15 | 2.36 | 1.24 | 0.396 | 0.436 | 0.538 |

|  |  |  |  |  |  |  |  |  |
| --- | --- | --- | --- | --- | --- | --- | --- | --- |
| *Stenotrophomonas* | 1.52 | 0.23 | 4.16 | 0.01 | 2.27 | 0.600 | 0.242 | 0.536 |
| *Enterococcus* | 1.40 | 3.14 | 0.52 | 1.35 | 1.33 | 0.323 | 0.342 | 0.735 |
| *Lactobacillus* | 0.01 | 4.06 | 0.00 | 2.41 | 1.89 | 0.664 | 0.100 | 0.669 |

## DISCUSSION

New data obtained from both human (He et al., 2020; Rackaityte et al., 2020) and bovine fetuses (Guzman et al., 2020; Husso et al., 2021) suggest that microbial colonization of the fetal gastrointestinal tract may take place before birth. In these studies, *in utero* microbial colonization was investigated during mid or late gestation. In the present study, we investigated the presence of a microbial community in bovine fetuses during early gestation. For this, we characterized microbiota associated with amniotic and allantoic fluids as well as intestinal and placental tissues in 83-day-old calf fetuses, which represents the end of the first trimester of pregnancy. Our 16S rRNA gene sequencing, qPCR (intestinal and placental tissues) and SEM imaging (fetal fluids) data derived from four fetal compartments and, most importantly, from the fetuses with the same sex (female) and same gestational age (83-day-old) and born from the dams that were genetically homogenous and managed in the same farm provide evidence for the presence of bacterial microbial communities that are being established in bovine fetuses during the first trimester of gestation.

The fetal bacterial microbiota was dominated by the Proteobacteria, Firmicutes, Actinobacteriota and Bacteroidota phyla. Similar to our study, previous studies reported the presence of an archaeal and bacterial microbiota in amniotic fluid and the intestine of 5-, 6- and 7-month-old fetuses (Angus × Friesian bred) (Guzman et al., 2020), and C-section derived near full-term calf fetuses (Belgian Blue) (Husso et al., 2021). The dominant bacterial phyla observed in the 83-day-old fetuses in the present study were also the predominant phyla in these mid- and full-term calf fetuses and in the calves postnatally. These phyla are key members of the bovine respiratory (Timsit et al., 2020), gastrointestinal (Holman and Gzyl, 2019) and reproductive tract (Amat et al., 2021), as well as the mammary gland microbiota (Khafipour, 2016; Derakhshani et al., 2018). Thus, the bacterial community profile in the 83-day-old fetuses resembles that of the fetus during mid- or late-gestation and after birth at the phylum level.

Such similarity in bacterial composition between 83-day-old fetuses and mid- or late-term calf fetuses (Guzman et al., 2020; Husso et al., 2021) was also partially observed at higher taxonomic resolution, with eight out of the 20 most relatively abundant genera in the 83-day-old fetuses (*Acinetobacter, Anoxybacillus, Corynebacterium, Enterococcus, Sphingomonas, Staphylococcus, Stenotrophomonas* and *Streptococcus*) also being dominant in both mid and full-term fetuses. Bacterial genera such as *Flavobacterium, Bradyrhizobium, Enhydrobacter, Sphingomonas, Pseudomonas* and *Lactobacillus* that are commensals found in raw cow’s milk and colostrum (Rainard, 2017; Klein-Jöbstl et al., 2019; Taponen et al., 2019) were also identified as dominant genera here. In addition to these genera, *Brucella* and *Saccharopolyspora*, which encompass pathogenic species, were also dominant in the 83-day-old fetuses. *Brucella abortus* is often isolated from aborted bovine fetuses (López et al., 1984; Xavier et al., 2009) and *Saccharopolyspora rectivirgula* (formerly known as *Micropolyspora faeni*) can induce lung infections in both cattle and cattle farmers if inhaled (Gershwin et al., 1994).

Apart from those bovine host-specific bacterial species, non-host bacterial species including ANPR (present in corn silage) (Wang et al., 2021) and *Aphanizomenon* (filamentous nitrogen-fixing cyanobacteria which can produce toxic metabolites such as neurotoxins and cytotoxins) (Cirés and Ballot, 2016) were also detected in high abundance in fetal fluids. Given that the fetal fluids were obtained from the intact fetus using a sterile needle and syringe, and that all OTUs found in the negative extraction controls were removed, it is less likely that these non-host specific taxa found in amniotic and allantoic fluid samples may be contaminants.

When compared among different fetal sample types, bacterial community structure was distinctly different between the amniotic and allantoic fluids, and the fetal intestine and placenta of 83-day-old calf fetuses. Fetal fluids also had a more diverse and richer bacterial community compared to the intestinal and placental tissues. This observation is consistent with previous studies where microbial diversity in differed between amniotic fluid and the gastrointestinal tract of mid- and full-term calf fetuses (Guzman et al., 2020; Husso et al., 2021). The four fetal sample types also differed here in terms of phyla with two of the eight most predominant phyla (Proteobacteria and Acidobacteriota), and six out of the 20 most relatively abundant genera, differing in relative abundance between the different fetal sample types. At higher taxonomic resolution each of the fetal compartments harbored a number of unique bacterial taxa, with only 55 OTUs found in at least one sample from each of the four fetal tissue/fluid types (Fig. 3 and 4). This indicates that a distinct bacterial microbiota may be present in fetal fluids, and the intestine and placenta during early gestation. Although many factors may be responsible for the distinct microbial communities within these fetal compartments, the physiological, biochemical and immunological properties that are unique to amniotic and allantoic fluids, and the mucosal surfaces of intestine and placenta may play a major role in shaping these microbiota (Underwood et al., 2005; Mei et al., 2019; Stras et al., 2019; Husso et al., 2021).

There was also considerable variation in terms of taxon prevalence and relative abundance between individual fetuses within each sample type. This was somewhat unexpected considering the common gestational age- (83 days-old), sex- (female) and maternal-homogeneity of the fetuses used in this study. Indeed, greater individual variation in the fetal microbiota was previously observed in calf fetuses (Guzman et al., 2020; Husso et al., 2021). In the study by Husso et al., (2021), 25 full term calf fetuses were obtained via C-section over a 16-month period and 7 of these fetuses were male. The calf fetuses used by Gusman et al., (2020) were 5, 6 and 7 months old, with unreported sexes, gestational ages, and sources and genetic backgrounds of the dams. Guzman et al. (2020) observed that total bacterial abundance (determined by qPCR) in amniotic fluid and intestine of mid-term fetuses increased with gestational age. Thus, besides gestational age, it is plausible that differences in fetal sex and heterogeneity associated with the mother’s genetic background, diet and management practices could led to inter-fetus variations in the microbiota. The low bacterial biomass present in the fetal samples, which was evident by the qPCR-estimated total bacterial concentration in the fetal intestine and placenta tissues (Fig.5A), is likely another factor contributing to the greater inter-individual variation in the fetal-associated microbiota.

Another potential factor that could influence the fetal microbiota is the maternal diet. Although the impact of maternal nutrition in fetal programming has been relatively well documented in both humans and cattle (Palmer, 2011; Caton et al., 2019), and diet-based alteration of the maternal gut microbiota during gestation has been shown to modulate fetal metabolic and neurodevelopment in mice (Kimura et al., 2020; Vuong et al., 2020), the impact of maternal nutrition on *in utero* microbial colonization remains relatively unexplored. Nutrition is a key factor that influences the maternal gastrointestinal (Carvalho et al., 2020), and reproductive microbiota (Xu et al., 2021) and both are considered to be main inoculum sources for the uterine microbiota (Altmäe, 2018; Baker et al., 2018). Therefore, it has been hypothesized that maternal nutrition may also influence *in utero* microbial transmission. Our results from the fetuses whose mothers were subjected to different rates of gain and diets supplemented with or without VTM supplementation suggest that alteration of maternal diet may influence fetal microbial colonization during early gestation. Although the microbial community structure and diversity did not differ by maternal treatment in any of the fetal sample types, difference in microbial richness of allantoic fluid and intestinal microbiota, and abundance of some predominant phyla (Proteobacteria in amniotic fluid and the intestine, Firmicutes in the placenta, and Cyanobacteria and Bacteroidota in amniotic fluid) was detected between maternal diet treatment groups.

Some of the potential limitations of the present study include: 1) small sample size (n = 8), and 2) unavailability of data from the dams of these fetuses showing the impact of the dietary interventions on the maternal microbiota during pregnancy and/or at the time of fetal tissue harvesting. As a result, it is difficult to make a definitive conclusion regarding the impact of maternal rate of gain and VTM supplementation on the bovine fetal microbial profile during early gestation. Nevertheless, the results of this study provide, for the first time, 16S rRNA gene sequencing-based evidence indicating that maternal nutrition may influence *in utero* microbial colonization in cattle fetuses. Future studies with a larger sample size, and longitudinal samples from the maternal gut and reproductive microbiota in response to diet alteration is warranted to confirm the impact of maternal nutrition during pregnancy on fetal microbial colonization. In addition to sequencing methods, culturing should be included to isolate viable bacteria present in the fetal intestine and fluids. Identifying the role of the bovine maternal microbiota during pregnancy in the development of the fetal microbiome, as well as offspring metabolic and microbial programming has important implications for the cattle industry. Maternal microbiome-targeted approaches to influence perinatal microbial colonization may lead to novel ways to enhance feed efficiency and disease resilience in cattle.

In summary, our results revealed the presence of a relatively diverse and complex bacterial community in allantoic and amniotic fluids, and fetal intestine and placenta on day 83 of gestation in beef cattle. Overall, the fetal bacterial community was dominated by Proteobacteria, Firmicutes, Actinobacteriota and Bacteroidetes. Microbial community structure and diversity was significantly different between allantoic and amniotic fluids, and the fetal intestine and placenta. Total bacterial load in the intestinal and placental samples as well as SEM imaging of the fetal fluids provided additional evidence for the presence of a microbiota in these samples. Furthermore, minor effects of maternal rate of gain and dietary VTM supplementation on the fetal microbiota were detected. Overall, the results of this study, for the first time, indicate that colonization with pioneer microbiota may occur during the first trimester of gestation in bovine fetuses, and that the maternal nutritional regime during gestation may influence the fetal microbiota.

## Authors’ contributions

Conceiving the idea, designing the study, and providing supervision—S.A., C.R.D.; Cattle management— K.L.M, C.R.D. and K.K.S.; Animal care and sample collections—C.R.D, K.L.M, S.T.D, A.K.W., P.P.B, L. P. R, J. S. C; Sample processing—K.S. and S.A.; Bioinformatics analysis— D.B.H. and S.A.; Data processing and statistical analysis—D.B.H and S.A.; Manuscript writing—S.A.; Manuscript review and editing—S.A., D.B.H., C.R.D, J.S.C., A.K.W and L. P. R,, All authors have read and agreed to the published version of the manuscript.

## Figure legends

**Supplementary Figure S1.**
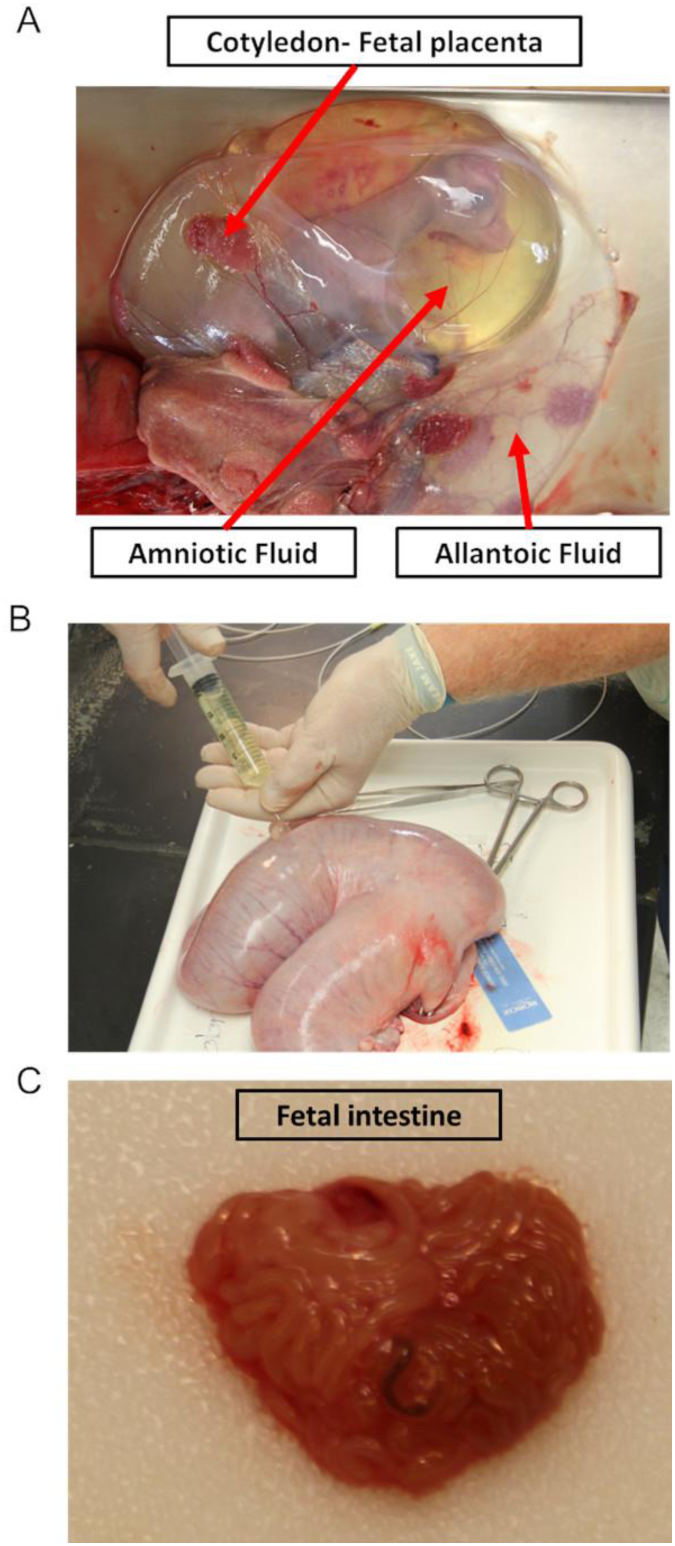
**A**) The anatomic location of amniotic and allantoic fluid, and fetal placenta (cotyledon), **B**) the collection of the fetal fluids, and **C**) the fetal intestine of the 83-day-old calf fetuses.

---

**Supplementary Table S1.**
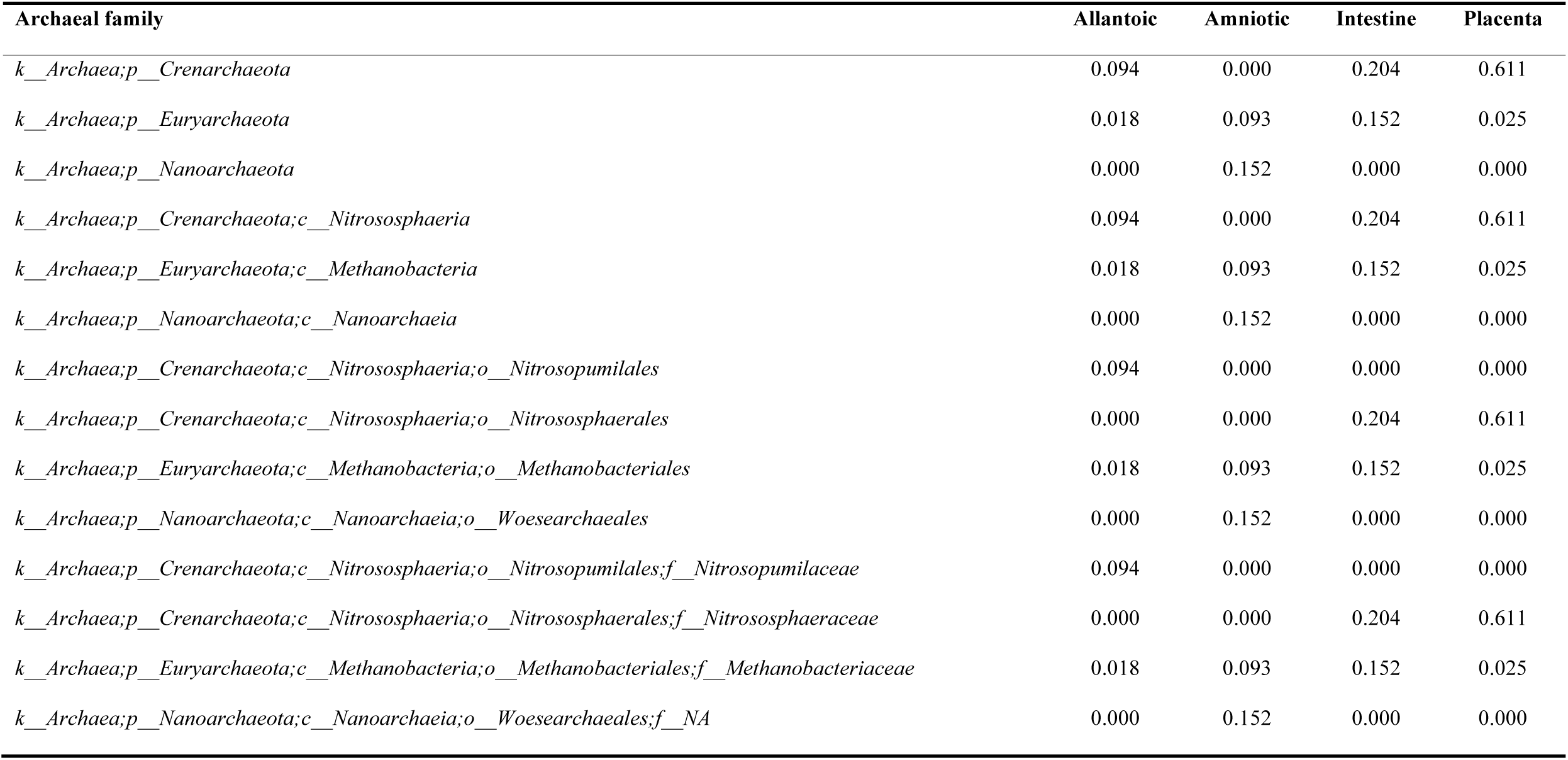
Overall relative abundance (%) of archaeal families present in allantoic and amniotic fluid, and intestinal and placental tissue samples obtained from 83-day-old calf fetuses.

| Archaeal family | Allantoic | Amniotic | Intestine | Placenta |
| --- | --- | --- | --- | --- |
| *k__Archaea;p__Crenarchaeota* | 0.094 | 0.000 | 0.204 | 0.611 |
| *k__Archaea;p__Euryarchaeota* | 0.018 | 0.093 | 0.152 | 0.025 |
| *k__Archaea;p__Nanoarchaeota* | 0.000 | 0.152 | 0.000 | 0.000 |
| *k__Archaea;p__Crenarchaeota;c__Nitrososphaeria* | 0.094 | 0.000 | 0.204 | 0.611 |
| *k__Archaea;p__Euryarchaeota;c__Methanobacteria* | 0.018 | 0.093 | 0.152 | 0.025 |
| *k__Archaea;p__Nanoarchaeota;c__Nanoarchaeia* | 0.000 | 0.152 | 0.000 | 0.000 |
| *k__Archaea;p__Crenarchaeota;c__Nitrososphaeria;o__Nitrosopumilales* | 0.094 | 0.000 | 0.000 | 0.000 |
| *k__Archaea;p__Crenarchaeota;c__Nitrososphaeria;o__Nitrososphaerales* | 0.000 | 0.000 | 0.204 | 0.611 |
| *k__Archaea;p__Euryarchaeota;c__Methanobacteria;o__Methanobacteriales* | 0.018 | 0.093 | 0.152 | 0.025 |
| *k__Archaea;p__Nanoarchaeota;c__Nanoarchaeia;o__Woesearchaeales* | 0.000 | 0.152 | 0.000 | 0.000 |
| *k__Archaea;p__Crenarchaeota;c__Nitrososphaeria;o__Nitrosopumilales;f__Nitrosopumilaceae* | 0.094 | 0.000 | 0.000 | 0.000 |
| *k__Archaea;p__Crenarchaeota;c__Nitrososphaeria;o__Nitrososphaerales;f__Nitrososphaeraceae* | 0.000 | 0.000 | 0.204 | 0.611 |
| *k__Archaea;p__Euryarchaeota;c__Methanobacteria;o__Methanobacteriales;f__Methanobacteriaceae* | 0.018 | 0.093 | 0.152 | 0.025 |
| *k__Archaea;p__Nanoarchaeota;c__Nanoarchaeia;o__Woesearchaeales;f__NA* | 0.000 | 0.152 | 0.000 | 0.000 |

**Supplementary Table S2.** *P-values* for the comparison in alpha diversity metrics and relative abundance of the five relatively most abundant phyla associated with different fetal samples between treatment groups.

| P-values | Allantoic |  |  | Amniotic |  |  | Intestine |  |  | Placanta |  |  |
| --- | --- | --- | --- | --- | --- | --- | --- | --- | --- | --- | --- | --- |
|  | Gain | VTM | Gain × VTM | Gain | VTM | Gain × VTM | Gain | VTM | Gain × VTM | Gain | VTM | Gain × VTM |
| ***Alpha diversity*** |  |  |  |  |  |  |  |  |  |  |  |  |
| Number of OTUs | 0.055 | 0.295 | 0.459 | 0.828 | 0.880 | 0.359 | 0.477 | 0.226 | 0.054 | 0.366 | 0.765 | 0.241 |
| Shannon diversity index | 0.302 | 0.311 | 0.565 | 0.654 | 0.772 | 0.122 | 0.723 | 0.805 | 0.100 | 0.626 | 0.942 | 0.740 |
| Inverse Simpson's index | 0.453 | 0.536 | 0.191 | 0.862 | 0.851 | 0.180 | 0.828 | 0.927 | 0.117 | 0.454 | 0.935 | 0.463 |
| ***Top 5 phyla*** |  |  |  |  |  |  |  |  |  |  |  |  |
| *Proteobacteria* | 0.920 | 0.872 | 0.743 | 0.188 | 0.448 | 0.040 | 0.163 | 0.002 | 0.000 | 0.755 | 0.610 | 0.630 |
| *Actinobacteriota* | 0.561 | 0.691 | 0.786 | 0.626 | 0.395 | 0.119 | 0.721 | 0.905 | 0.668 | 0.461 | 0.789 | 0.763 |
| *Firmicutes* | 0.760 | 0.735 | 0.149 | 0.640 | 0.732 | 0.919 | 0.645 | 0.006 | 0.343 | 0.168 | 0.066 | 0.692 |
| *Bacteroidota* | 0.760 | 0.735 | 0.149 | 0.057 | 0.363 | 0.173 | 0.554 | 0.741 | 0.308 | 0.934 | 0.198 | 0.806 |
| *Cyanobacteria* | 0.687 | 0.696 | 0.341 | 0.169 | 0.014 | 0.262 | NA | NA | NA | NA | NA | NA |
| *Acidobacteriota* | NA | NA | NA | NA | NA | NA | 0.125 | 0.766 | 0.123 | 0.241 | 0.636 | 0.082 |

